# Female song elicits increased vocal display effort in both sexes of Alston’s singing mouse

**DOI:** 10.1101/2024.11.19.624417

**Authors:** Joel A. Tripp, Steven M. Phelps

**Affiliations:** Department of Integrative Biology, University of Texas at Austin, Austin, TX; Program of Neuroscience, Carleton College, Northfield, MN

## Abstract

Acoustic displays are conspicuous behaviors common across diverse animal taxa. They have long been studied in behavioral ecology, evolutionary biology, and neuroscience. Most of these investigations, however, have focused on male display. For species in which both sexes display, correcting this bias will lead to a fuller understanding of the evolution and function of these behaviors. In this study, we investigated the role of vocal advertisement display in female Alston’s singing mice (*Scotinomys teguina*). Singing mice are small muroid rodents of the family Cricetidae, native to high altitude cloud forests of Central America. Females and males both produce elaborate advertisement songs used in mate attraction and intra-sexual competition. Prior studies have largely focused on male behavior, though we recently found that both sexes dramatically increase their rate of singing and song duration in response to playback of conspecific male song. Here, we tested how mice of both sexes adjust their song effort in response to female song playback. Our results show that, like male song, female song elicits robust increases in song effort from both female and male mice. This study reveals additional social contexts that promote high song effort in this species, while raising additional questions about the role of song in communication within and between the sexes.

---

Acoustic displays are among the most wide-spread and conspicuous signals across the animal kingdom. Despite this fact, the vast majority of research on acoustic display behaviors has focused either on male display or female responses to male display (e.g. Ballentine, 2004; Burkhard, Sachs, et al., 2023; Gustison & Bergman, 2016; Hovi et al., 1997; Mennill et al., 2002; Pasch, George, Campbell, et al., 2011; West & King, 1988). More recently, there is a growing understanding that female display is more common than previously appreciated (Odom et al., 2014). In order to fully understand the evolution and function of these behaviors, it is necessary to know how they are employed and how they affect members of both sexes.

The importance of understanding female display has been most strongly recognized by the songbird research community (Odom & Benedict, 2018). Over the past decade, that field has shown that female song is more common than once appreciated and is in fact the ancestral state (Odom et al., 2014). Like their male counter-parts, female songbirds use song to defend territories, attract mates, and coordinate reproduction (Langmore, 1998). Females also engage in duetting (Hall, 2009) and vocal mimicry (Dalziell & Welbergen, 2016; Gammon & Stracey, 2022). In mammals, female vocal displays have been found to function in territory defense (Cowlishaw, 1992; Petric & Kalcounis-Rueppell, 2013), maintaining bonds (Méndez-Cárdenas & Zimmermann, 2009), and signaling receptivity (Manno et al., 2008; Neunuebel et al., 2015). While the function of female displays will vary across species due to social and ecological selective forces, gaining a better understanding of these behaviors in females in addition to males will deepen our understanding of the evolution of display behaviors.

Alston’s singing mice (*Scotinomys teguina*) provide an ideal system for investigating acoustic display across the sexes. Named for their distinctive trilled vocalization, singing mice are native to the montane cloud forests of Central America (Hooper, 1972). Their song consists of rapidly repeated down-sweeping notes that begin in the ultrasonic and end in the human-audible frequency range (Fig. 1a). While both sexes sing, males tend to do so more frequently and produce longer songs on average than females (Miller & Engstrom, 2007), though there is significant overlap in song length across the sexes. Prior research has largely focused on male song, and has established that males increase their rate of singing following a brief interaction with a female (Fernández-Vargas et al., 2011) and in response to conspecific male song (Okobi et al., 2019; Pasch et al., 2013), indicating a role in both inter- and intrasexual communication.

**Figure 1.**
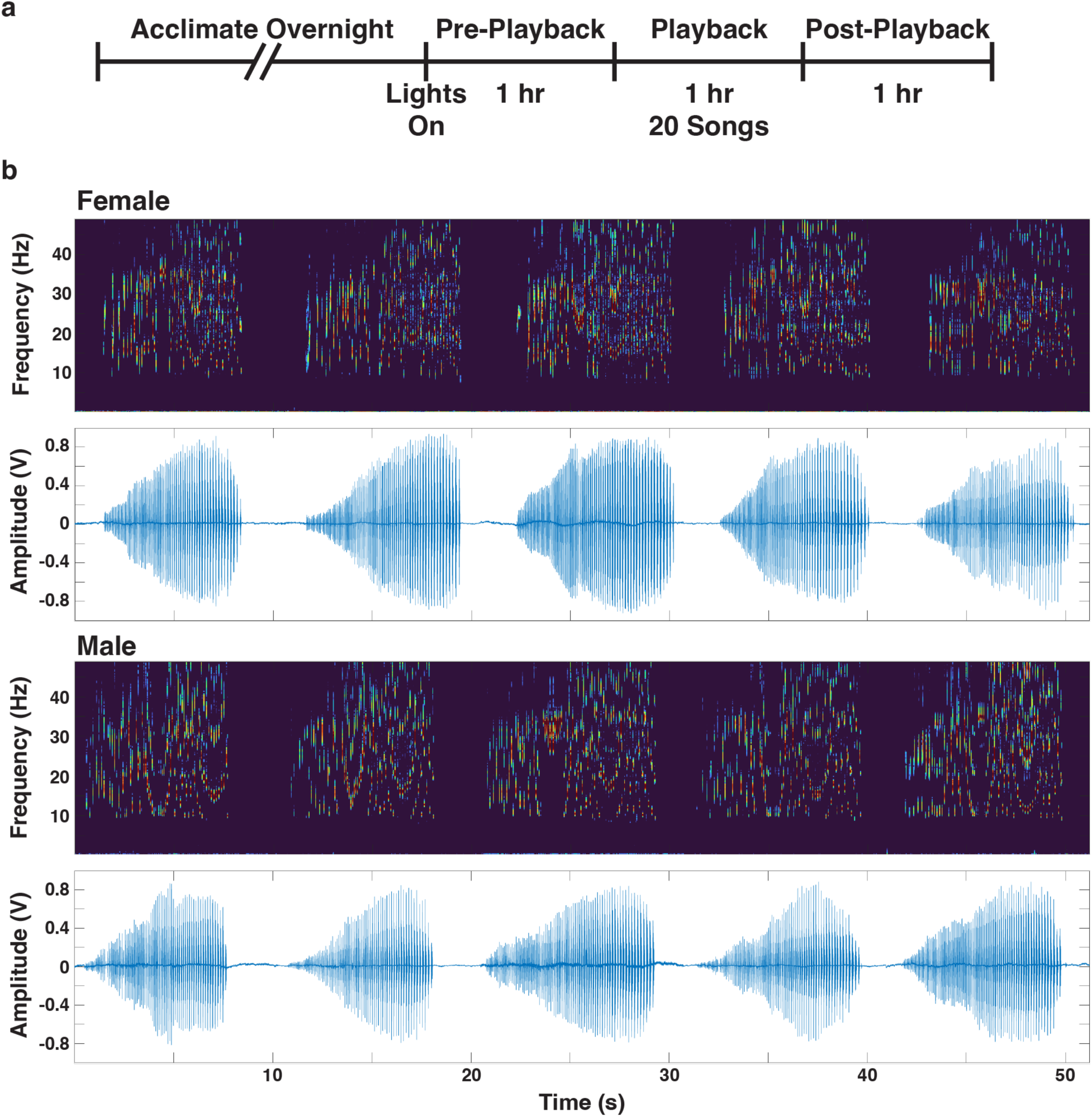
Experiment design and stimuli. a) Playback experiment timeline. For experiment comparing responses to female and male stimuli within subjects (Fig. 3), mice were kept in chambers and recorded on two consecutive days. b) Spectrograms and oscillograms of example female and male stimulus sets. Examples shown made up Set A (SGTf27 and SGTm100, see Table 2 for details) used in experiment comparing responses to female and male stimuli within subjects.

More recently (Tripp & Phelps, 2024), we demonstrated that although there are differences between the sexes, females also respond to male song playback by increasing the rate of singing, increasing song length, and producing counter-songs, in which an animal precisely times it’s song to begin with only a short latency (milliseconds to seconds) after the end of another animal’s song (Okobi et al., 2019). However, whether and how mice of either sex adjust their song effort in response to female song remains unknown. In this study, we first tested whether mice of either sex increase their song effort (i.e. song rate and duration) in response to female song playback and then sought to understand whether singing mice respond differently to female versus male song. Our results demonstrate that just like male song, female song is a salient social signal to both sexes, further elucidating the role of this vocal display.

## Methods

### Animal Subjects

Subjects were adult singing mice (*S. teguina*) of both sexes reared in a colony at UT Austin, derived from wild-caught animals captured in San Gerardo de Dota Valley, Costa Rica (n=12 per sex for experiment testing responses to female song playback and n=12 per sex for experiment comparing responses to female versus male song). Prior to playback experiments, subjects were housed singly or in groups of same-sex littermates in hamster cages equipped with running wheels, PVC tubing, nestlets, and sphagnum moss that was misted regularly with water to maintain cage humidity. Water and food (50% Lab Diet feline chow and 50% Lab Diet insectivore chow by weight) were provided *ad libitum*. Colony and testing rooms were maintained at 19-22° C on a 12 h:12 h light:dark cycle. Group housed animals were separated into individual home cages at least 24 h prior to testing.

### Playback experiment

Playback methods followed our previous study (Tripp & Phelps, 2024). Singly housed mice were transferred in their home cages to an acoustic isolation chamber and allowed to acclimate overnight. The following morning, at the beginning of the light period when mice are typically most vocally active, the acoustic chamber door was opened, and animals were recorded for three hours (Figure 1a). Acoustic recordings were made using a microphone and preamplifier (ACO Pacific, Belmont, CA) connected to an MA-3 amplifier and RX8 microprocessor (Tucker-Davis Technologies, Alachua, FL) at 32-bit resolution with a sampling rate of 97.7 kHz. During the second hour of recording, mice were presented with playback of conspecific songs, played from a speaker (Vifa, Viborg, DK) in an adjacent acoustic chamber. This approach has previously been found to be effective at consistently eliciting increased song effort and counter-songs in singing mice (Okobi et al., 2019; Tripp & Phelps, 2024). Recording and playback were controlled by custom Matlab (v9.7.0) and RPvdsEx (v92) scripts.

Song stimuli were recorded from wild-caught animals unfamiliar to experimental subjects (Tables 1 and 2). Each stimulus set consisted of five songs from a single individual (Figure 1b), and subjects received 20 playbacks from a single stimulus set at randomized time intervals over the playback hour, resulting in each song in the set being played four times. We first used playback of female stimuli to test whether female or male mice increase their song effort or counter-sing in response to female song. In this experiment, each animal was recorded for one day, then returned to the colony. In a second experiment, we compared changes in song effort and counter-singing of both sexes in response to both male and female song. For this experiment, the acoustic chamber door was closed following the first recording session, animals remained in the recording chamber overnight and were recorded and received playback following the same methods on the second day. The order of stimulus sex presentation was alternated across subjects. Three stimulus sets were used for testing male and female responses to female song playback and the same three sets were used along with three male stimulus sets for testing responses to female versus male stimuli (Table 2). For this experiment, female and male stimuli were recorded from animals in the same population (Burkhard, Matz, et al., 2023) and were matched to have the same mean song length. All song stimuli were normalized to a peak amplitude of 1 V and were played out at an amplitude of 4 V.

**Table 1.** Stimulus sets used to test response to either male or female song separately (related to Figure 2).

| Animal ID | Population | Figure | Sex | Mean Note Number | Mean Call Length (s) |
| --- | --- | --- | --- | --- | --- |
| CGm8 | Cerro Gomez | 2a-c | M | 77.2 | 7.78 |
| CGm21 | Cerro Gomez | 2a-c | M | 105.2 | 8.56 |
| CGm27 | Cerro Gomez | 2a-c | M | 94.8 | 7.86 |
| SGTf27 | San Gerardo de Dota | 2d-f | F | 99.8 | 8.17 |
| STGf28 | San Gerardo de Dota | 2d-f | F | 79.4 | 6.55 |
| SGTf33 | San Gerardo de Dota | 2d-f | F | 90.8 | 7.80 |

### Statistical analysis

Audio recording files were analyzed using custom Matlab scripts. Raw recordings were bandpass filtered at 5-48 kHz, then squared and down sampled to 100Hz using resample. Songs were identified as samples above an empirically determined threshold and were extracted for analyses of acoustic features as previously described (Burkhard et al., 2018; Campbell et al., 2010). One half second was used as the threshold to identify vocalization as separate songs. Counter-songs were defined as songs produced within 20 s of the onset of a song playback and if multiple songs were sung in this period, only the first was counted as a counter-song (Tripp & Phelps, 2024).

All statistical comparisons were made with R (v. 4.2.2) using the RStudio environment (v. 2023.06.0+421). Linear models were fit using lme4 (v. 1.1-31) (Bates et al., 2015) followed by ANOVA with type III sum of squares implemented by the car package to calculate F statistics and p values for each fixed effect (Fox & Weisberg, 2019). We used the emmeans package (Lenth et al., 2022) to perform *post hoc* t-tests and calculate p-values, which were Tukey-adjusted for multiple comparisons.

Song count data were highly skewed and were log2-transformed prior to analyses. For the experiment testing the effect of female song playback, transformed counts and mean song durations were fit to a linear mixed models with period, sex, and period x sex interaction as fixed effects, and subject as a random effect; counter-songs were compared across sex using a generalized linear model with a quasipoisson distribution. For the experiment comparing the effects of female and male song playback, transformed counts were fit to a linear mixed model with period, sex, period x sex interaction, and stimulus sex as fixed effects and playback day and subject as random effects. Counter-songs were compared across sex and stimulus sex using a generalized linear model with quasipoisson distribution.

### Ethical Note

All procedures were approved by the Institutional Animal Care and Use Committee of the University of Texas at Austin and followed guidelines set by the National Institutes of Health Guide for the Care and Use of Laboratory Animals. When feasible, animals were returned to the colony following these studies for use in future experiments. Animals that could not be used in further studies were euthanized by carbon dioxide inhalation followed by decapitation in accordance with institutional guidelines

## Results

### Female song elicits increased song effort from both sexes

Previously, we found that both sexes of singing mice increase their song effort in response to playback of conspecific male song (Fig. 2a-c), with males showing greater increases in song rate and length than females (Tripp & Phelps, 2024). Here, we tested the effects of female song playback across both sexes. We found that, as with male song stimuli, both female and male singing mice significantly increase their rates of singing in response to conspecific female song; however, unlike the response to male song, we found no sex differences in the rate of singing in response to female song playback (Fig. 2d). A linear mixed model predicting the song rate across each recording period by sex showed a significant effect of period (F(2,44) = 16.22, p = 5.29*10^-6^), but not sex (F(1, 28.9) = 0.0048, p = 0.95), nor a period x sex interaction (F(2, 44) = 1.21, p = 0.31). Pairwise comparisons found that both sexes sang significantly more during the playback hour compared to pre-playback (Females: t44 = 5.22, p < 0.0001; Males: t44 = 3.12, p = 0.0088) and post-playback hours (Females: t44 = 4.59, p = 0.0001; Males: t44 = 4.12, p = 0.0005).

**Figure 2.**
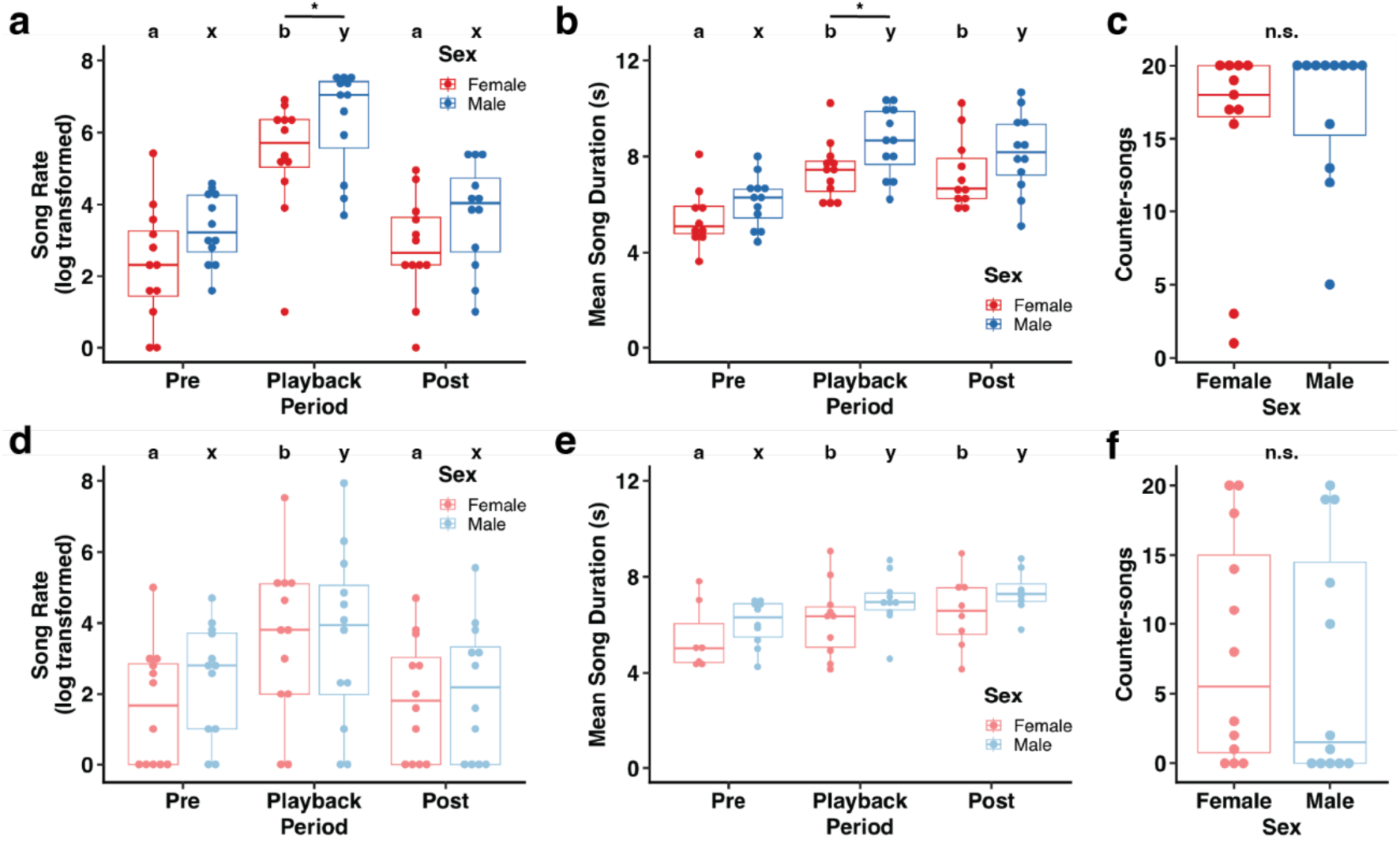
Male and female song playback elicits increased song effort. a) Female and male singing mice increase their song rates in response to playback of male song, though males sing more than females during the playback period. b) Females and males both increase their mean song length during and after playback, compared to the pre-playback period. Male songs are significantly longer than females’ during the playback period. c) Females and males counter-sing at similar rates in response to male song playback. d) Both sexes increase song rates in response to female song playback, there are no differences in song rate across the sexes. e) Both sexes increase mean song length during the playback and post-playback period compared to pre-playback, there are no differences in song length across the sexes. f) Females and males counter-sing at similar rates in response to female song playback. Letters indicate significant differences within sex across periods, asterisks indicate differences across sexes within periods. Data presented in a-c previously published (Tripp & Phelps, 2024).

Similarly, we found that both sexes sing longer songs during the female song playback period and post-playback compared to the pre-playback period (Fig. 2e). As with song rate, there was no difference in song length across the sexes in this experiment. A linear mixed model predicting song length across playback period and sex showed a significant effect of period (F(2, 28.6) = 15.98, p = 2.19*10^-5^), but not sex (F(1, 23.1) = 1.12, p = 0.30), nor a period x sex interaction (F(2, 28.6) = 0.77, p = 0.47). Pairwise comparisons showed that females and males each sang shorter songs during the pre-playback hour compared to the playback (Females: t28.7 = 5.01, p = 0.0001; Males: t28.9 = 3.74, p = 0.0023) and post-playback (Females: t28.6 = 5.02, p = 0.0001; Males: t28.6 = 4.21, p = 0.0007) hours. Rates of counter-singing did not differ between females and males (Fig. 2f; F1 = 0.10, p = 0.76).

### Stimulus song sex does not impact song effort

Surprisingly, the sex differences we observed in song rate and length in response to male song were absent when singing mice were presented with conspecific female song playback. Additionally, female song playback tended to elicit lower song rates than male song playback (Fig. 2). Due to these observations, we next tested how individual singing mice responded to playback of both male and female songs.

We once again found that playback of songs from either sex elicits increased song rates in both female and male mice (Figure 3a). Our results also showed that males had higher rates of singing than females at baseline, but not during song playback. A linear mixed model predicting song rate across each recording period by subject sex and stimulus sex showed significant effects of period (F2,114.0 = 28.4, p = 9.81*10^-11^), sex (F1,29.3 = 7.29, p = 0.011), and period x sex interaction (F2,114.0 = 3.67, p = 0.028), but not stimulus sex (F1,114.0 = 0.54, p = 0.46). *Post hoc* pairwise comparisons showed that both sexes sang more often during the playback period compared to pre-playback (Females: t114 = 6.81, p < 0.0001; Males: t114 = 2.98, p = 0.0099) and post-playback (Females: t114 = 6.203, p <0.0001; Males: t114 = 4.31, p = 0.0001). Additionally, males sang more songs than females during the pre-playback period (t29.3 = 2.70, p = 0.011). We also noticed a trend toward higher song rates in males post-playback (t29.3 = 1.83, p = 0.078); however, both sexes sang at similar rates during playback (t29.3 = 0.97, p = 0.34).

**Figure 3.**
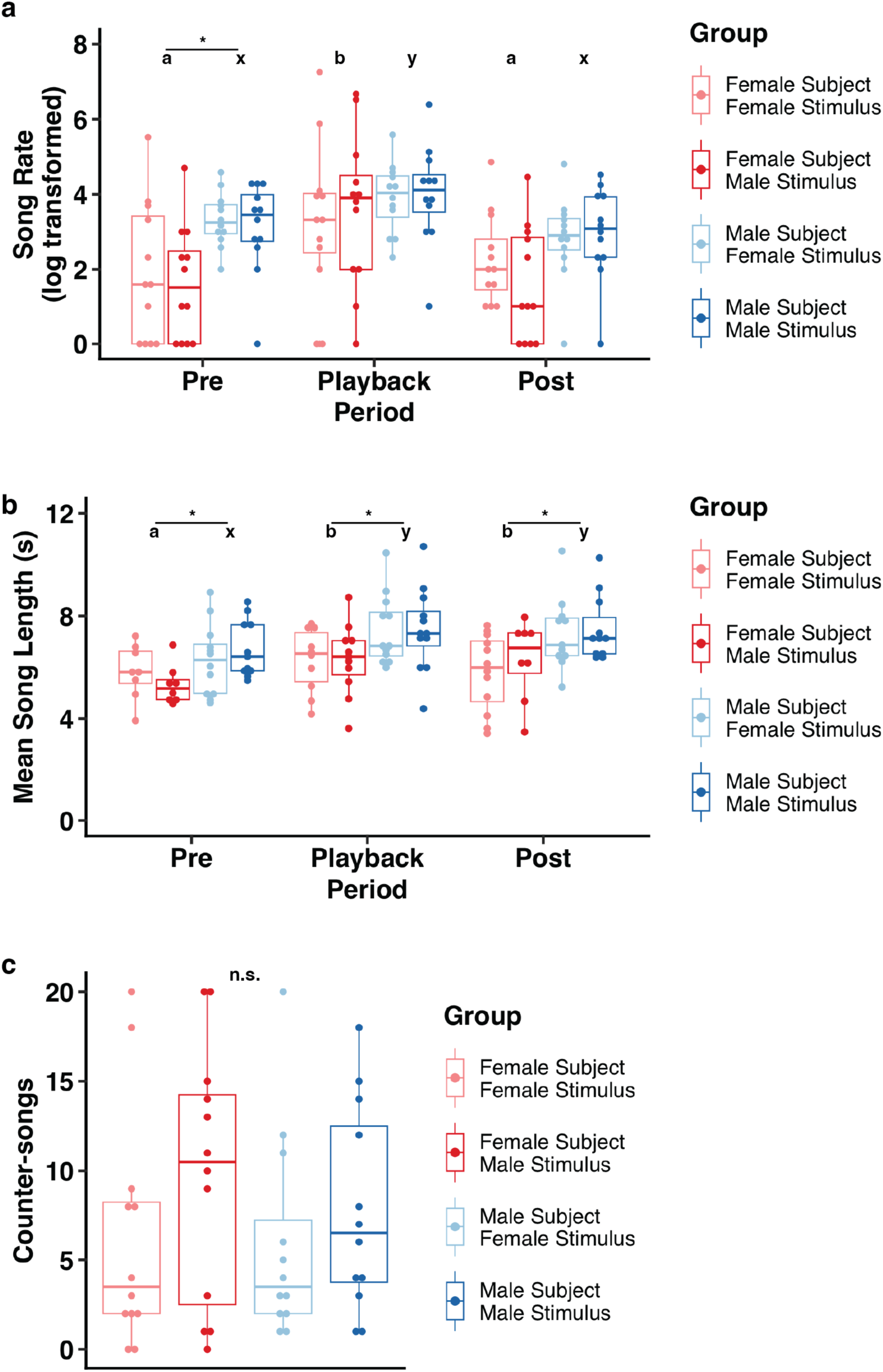
Song effort is not influenced by stimulus song sex. a) Both sexes increase song rates in response to playback at similar levels regardless of stimulus animal sex. b) Females and males both increase mean song lengths during playback and post-playback compared to the pre-playback hour. Males sing longer songs than females, but song length was not influenced by stimulus sex. c) Rate of counter-singing did not significantly differ across females and males or by stimulus sex. Letters indicate significant differences within sex across periods, asterisks indicate differences across sexes within periods.

We also found that song playback results in increased song length, regardless of stimulus song sex (Figure 3b). A linear mixed model predicting mean song length across each period by subject sex and stimulus sex showed significant effects of period (F2,97.6 = 14.8, p = 2.45*10^-6^) and sex (F1,27.7 = 7.47, p = 0.011), but not stimulus sex (F1,97.4 = 1.02, p = 0.31) or a period x sex interaction (F2,97.0 = 0.10, p = 0.90). Pairwise comparisons found that both sexes sang longer song during playback (Females: t97.3 = 5.12, p < 0.0001; Males: t96.1 = 5.35, p < 0.0001) and post-playback (Females: t97.6 = 4.43, p = 0.0001; Males: t96.4 = 4.61, p < 0.0001) compared to pre-playback. There were no differences between song lengths during the playback and post-playback periods (Females: t97.7 = 0.67, p = 0.78; Males: t96.3 = 0.58, p = 0.83). Across the sexes, we found that males sang longer songs than females during each recording period (pre-playback: t27.7 = 2.73, p = 0.011; playback: t25.9 = 2.55, p = 0.017; post-playback: t26.3 = 2.59, p = 0.015). We did not find significant differences in the rate of counter-singing based on focal animal sex (F1 = 0.47, p = 0.50) or stimulus sex (F1 = 2.14, p = 0.15)

## Discussion

In order to fully understand the evolution and function of display behaviors, it is necessary to know how they are employed and responded to by senders and receivers of both sexes. However, the role of female display has been understudied. Correcting this bias will provide deeper insight into the role of acoustic display behaviors in communication within and across the sexes.

Alston’s singing mice make excellent models for investigating these questions because both females and males readily display, but there are consistent differences across the sexes, suggesting variation in the function or value of display investment (Miller & Engstrom, 2007; Tripp & Phelps, 2024). In this study, we investigated responses to female song playback by both female and male singing mice. We found that playback of female song promotes higher song effort, as measured by singing rate and song length, in both sexes of singing mice. These results mirror previous findings on the effect of male song playback on singing mouse song effort.

Our previous study demonstrated that both sexes increase the rate of song production and length of their songs in response to playback of conspecific male song (Tripp & Phelps, 2024). Additionally, females and males counter-sang to conspecific male songs at similar rates. However, while both sexes increased song effort in response to male song, male effort was higher than that of females during playback. These results supported the previously identified role of song in intrasexual signaling for males (Okobi et al., 2019; Pasch et al., 2013; Pasch, George, Hamlin, et al., 2011) and suggested a role in intersexual signaling for females. Further, the results led us to question whether differences in effort observed across the sexes were due to stimulus sex. Thus, we sought to test whether either or both sexes responded to playback of female song with increased song effort and whether females would show higher effort than males when responding to a same-sex stimulus.

In the present study, we found that playback of female songs resulted in higher rates of singing and longer songs from both sexes compared to baseline behavior prior to playback. In addition, we observed no significant differences in the rate of counter-singing to female song between female and male subjects. However, unlike male stimulus songs, responses to female songs did not differ by sex in either rate or duration. Further, we noted that subject song rates tended to be lower in response to female song than we had previously observed in response to male song. To further investigate this difference, we next sought to directly compare responses to both female and male song within individual subjects of both sexes.

In our second experiment, we found that females and males significantly increased their song rates during playback of both female and male songs. Consistent with earlier observations (Miller & Engstrom, 2007; Tripp & Phelps, 2024), males tended to sing more than females, in this case before, but not during playback. Further, song playback also elicited increased song length from both sexes, with males singing longer songs than females within and outside playback periods. Surprisingly, however, we did not observe any effect of stimulus sex on song rate, length, or counter-singing. This result suggests that singing mice either do not adjust their song effort based on the sex of conspecifics they communicate with, or they are unable to determine the sex of stimulus animals based on song alone. This possibility is supported by the fact that aside from length, no consistent sex differences in song characteristics have been identified in singing mice (Burkhard, Matz, et al., 2023; Campbell et al., 2014; Miller & Engstrom, 2007). Mice may infer singer sex based on length; however song length distributions largely overlap (see Table 1), and we previously found no differences in response to stimulus sets of varying lengths (Tripp & Phelps, 2024). Further, male mice sing significantly more songs when given visual and olfactory access to a female compared to a male (Giglio, 2020), indicating that conspecific sex is likely a potent modulator of song effort. Still, it remains possible that the mice are capable of detecting differences that are not evident to researchers (Mulder et al., 2003), but respond with similar effort to both female and male singers.

One alternative is that the sex of a conspecific singer does influence song effort, but the association between a specific song and the singer’s sex must be learned. Singing mice exhibit consistent, heritable individual differences in the spectral features of their calls (Burkhard, Matz, et al., 2023) and have overlapping home ranges (Ribble & Rathbun, 2018), suggesting that mice may be capable of linking a specific song to a familiar conspecific through repeated exposure. This possibility is supported by evidence of vocalizations carrying information of identity in a number of avian (Briefer et al., 2013; Favaro et al., 2015; Gentner & Hulse, 1998; Mathevon, 1997; Pardo et al., 2018; Smith-Vidaurre et al., 2023) and mammalian species (Fukushima et al., 2015; Rubow et al., 2017; Schehka & Zimmermann, 2009; Toth & Parsons, 2018; Wierucka & Leu, 2024), including rodents (Blumstein & Munos, 2005; Vielle et al., 2021). Further, *S. teguina* males learn to avoid singing in response to songs produced by larger, more aggressive sister species, *S. xerampelinus*, in regions where they co-occur (Pasch et al., 2013), demonstrating they are capable of associating the acoustic features of a vocalization with prior social experiences. If this is indeed the case, singing mice may adjust their effort based on previous experience with a given individual. For example, a male may sing with higher effort in response to a song produced by a familiar female, representing a mating opportunity, but may sing less in response to a familiar aggressive male.

Somewhat surprisingly, we found minor variations in singing responses across each of our playback experiments. Previously, we found differences across the sexes in both song rate and length in response to male song playback, with males showing higher effort than females (Tripp & Phelps, 2024). In the current study, we found no differences in song rate during female song playback and only saw differences in length when including both male and female stimuli. Additionally, unlike our previous study focused on male song playback, the current experiments found sex differences in song rate and length outside of the playback period. These differences, especially those present during pre-playback, are unlikely to be caused by stimulus animal sex and in total, our results are consistent with a general trend that both sexes sing readily, but on average males invest more in singing than females, both in song rate and in song length (Miller & Engstrom, 2007). While these sex differences are consistent, they are also relatively low in magnitude and may or may not reach statistical significance in any given experiment.

Alternatively, it is possible that our results are reflective of real differences. One potential explanation is the use of stimulus songs from different populations between the current studies and previous experiment (Tripp & Phelps, 2024), as populations vary in measures of both effort and frequency modulation (Campbell et al., 2010). Additionally, the songs used for the male playback-only experiment are longer than the songs used for female playback and comparison of female and male playback by approximately 0.5 s on average. While we have not previously observed effect of playback song length on subject song effort (Tripp & Phelps, 2024), it remains possible that longer playback stimuli may influence males and females differently. The potential influence of stimulus song duration and source population led us to control for these factors in our experiment directly comparing the effect of female and male song playback (Table 2, Fig. 3). Still, neither of these factors is likely to influence the subjects’ behavior during the pre-playback period, in which we saw variation in sex differences in song effort across our experiments.

**Table 2.** Stimulus sets used to directly compare responses to female and male stimulus songs (related to Figure 3).

| Animal ID | Population | Set | Sex | Mean Note Number | Mean Call Length (s) |
| --- | --- | --- | --- | --- | --- |
| SGTf27 | San Gerardo de Dota | A | F | 99.8 | 8.17 |
| SGTm100 | San Gerardo de Dota | A | M | 99 | 8.11 |
| STGf28 | San Gerardo de Dota | B | F | 79.4 | 6.55 |
| SGTm117 | San Gerardo de Dota | B | M | 78 | 6.68 |
| SGTf33 | San Gerardo de Dota | C | F | 90.8 | 7.80 |
| SGTm129 | San Gerardo de Dota | C | M | 90.2 | 7.69 |

The results of the current study show that female song is an important social cue for singing mice of both sexes. Male song plays a role in both inter- and intrasexual signaling, likely important for both mate attraction and male-male competition (Burkhard, Sachs, et al., 2023; Fernández-Vargas et al., 2011; Pasch et al., 2013) and female song likely has a similar function, as both sexes robustly increase their song effort in response to playback, regardless of stimulus animal sex. Male song effort is positively associated with measures of body condition (Burkhard et al., 2018) and females show preference for higher effort songs (Burkhard, Sachs, et al., 2023; Pasch, George, Campbell, et al., 2011), indicating a role in signaling quality. In females, intersexual communication may be more focused on signaling receptivity, as in rats, mice and ground squirrels (Börner et al., 2016; Manno et al., 2008; Neunuebel et al., 2015; Warren et al., 2020; White & Barfield, 1987). The difference between signaling quality and receptivity may explain why males generally invest more in song effort than females.

As in males, female song is likely also involved in intrasexual competition. One small tracking study found that singing mice of both sexes had highly overlapping home ranges, though just two individuals of each sex were observed (Ribble & Rathbun, 2018). Another study, investigating a congener, the long-tailed singing mouse (*S. xerampelinus*) found that male ranges overlapped while females maintained more exclusive territories (Blondel et al., 2009). In the lab, males demonstrated higher intrasexual aggression and females showed signs of mutual avoidance (Blondel, 2006). This pattern is commonly seen in promiscuous rodents (Ribble & Stanley, 1998), and in one such species, the brush mouse (*Peromyscus boylii*), females have been found to use vocal interactions to mediate avoidance (Petric & Kalcounis-Rueppell, 2013), suggesting that female-female vocal interactions could be used as a means of avoiding one another or as a method of indirect competition. This possibility may explain why females increase song effort in response to playback.

One important caveat to our results is the potential role of estrus cycle in influencing female singing mouse behavior. We did not control or track estrus state in female mice in our playback studies, and so it remains possible that reproductive state influenced female behavior. This could be particularly important if female song functions as a signal of receptivity. However, prior evidence of male singing mouse responses to changes in female estrus cycles are mixed. While males spend more time investigating odors from females in estrus compared to non-estrus females (Fernández-Vargas et al., 2008), they do not modulate their vocal effort in response to female estrus state (Fernández-Vargas et al., 2011). Additional studies tracking how females change their investment in advertisement display may further elucidate the function of female song in signaling receptivity.

Beyond informing our understanding of the function of female singing mouse vocalization, this study adds to insights on the evolution of elaborate female display. While the reasons these displays evolve remains an open question, some patterns are emerging. In songbirds, several trends have been identified related to the retention of ancestral female song (Odom et al., 2014). Among these are tropical distributions (Price et al., 2009; Slater & Mann, 2004) and non-migratory behavior associated with year-round territoriality (Price, 2009; Robinson, 1949). Given their Neotropical distribution (Hooper, 1972) singing mice fit in with the first of these themes well. However, as discussed above, the association with territoriality is less clear. Lastly, while singing mice may not be fiercely territorial, they are not migratory, suggesting at least a partial parallel with another factor associated with the maintenance of female bird song.

In summary, our results show that playbacks of songs from either sex elicit increased song effort from both male and female singing mice. While we found differences in song effort across the sexes, these were not related to stimulus song sex, demonstrating that singing mice either do not adjust their song effort in response to stimulus animal sex or are unable to identify singer sex based on song alone. These results are consistent with a role for female song in signaling receptivity to males as well as intrasexual communication associated with competition or avoidance of other females and leave open the possibility that individual mice associate conspecific calls with specific individuals as potential mates or rivals.

## Acknowledgements

This work was supported by the U.S. National Institutes of Health (R01 NS113071 (S.M.P.) and F32MH125562 (J.A.T.)).

## Notes

### Competing Interest Statement

The authors have declared no competing interest.

